# Resident cardiac macrophages are not required for normal atrioventricular node conduction

**DOI:** 10.64898/2026.02.11.704506

**Authors:** Sami Al-Othman, Yakun Wu, Pierre Fontanaud, Franz Puttur, David Conesa, Chelsea Zhu, Sacha Moore, Roman Tikhomirov, Alice Francis, Sooraj Nair, Rasheda A Chowdhury, Mubassher Husain, Joseph J Boyle, Delvac Oceandy, Steven A Niederer, Richard Walton, Gareth Howell, Luke Roberts, Mark R Boyett, Michael A Colman, Matteo E Mangoni, Alicia D’Souza

## Abstract

Resident cardiac macrophages are understood to facilitate atrioventricular (AV) node conduction because they purportedly couple to AV node myocytes via connexin43 (Cx43) containing gap junctions. We tested this mechanism using biophysical modelling, high-resolution imaging of mouse and human AV conduction tissue, and pharmacological macrophage depletion. *In silico*, coupling macrophage membrane phenotypes to HCN4^+^ AV node myocytes imposed an electrotonic load that suppressed pacemaking and promoted conduction slowing, including stable 2:1 block in strand simulations. Anatomically, HCN4-defined components of the mouse AV conduction axis were essentially devoid of Cx43 and overlap of CD68+ macrophages and Cx43 was negligible in both mouse AV node and human penetrating bundle. Finally, near-complete macrophage depletion with CSF1R inhibition (PLX5622) did not alter AV electrical activity *in vivo* or *ex vivo*. Together, these data argue against a physiologically relevant role for Cx43-mediated macrophage–myocyte electrical coupling in normal AV node function.

**HIGHLIGHTS:**

- Modelling predicts that AV node automaticity and conduction would be suppressed if macrophages coupled to AV node myocytes
- The mouse AV conduction axis is essentially devoid of Cx43, currently considered responsible for macrophage-AV node myocyte coupling
- Overlap of macrophages and Cx43 expression is not discernible in the Cx43-expressing human distal AV node
- Macrophage depletion by CSF1R inhibition does not impact AV electrical activity *in vivo* or *ex vivo*

## INTRODUCTION

The atrioventricular (AV) node is a specialised component of the cardiac conduction system that delays ventricular excitation to allow filling of the ventricles, protects the ventricles from rapid atrial rhythms, and acts as a back-up pacemaker.^1,2^ These functions depend on the AV node’s distinctive electrophysiological properties and molecular makeup: slow diastolic depolarisation supported by HCN4^3^ and the L-type Cav1.3^4^ and T-type Ca_v_3.1Ca^2+^ channels,^5,4^ and importantly slow action potential conduction as a result of a slow L-type Cav1.2 channel-dependent action potential upstroke^4,6^ and weak intercellular coupling mediated predominantly by connexin 45^7,8^ rather than connexin 43 (Cx43; that is responsible for fast conduction in the working myocardium^9^). Using the fractalkine receptor CX3CR1 as a macrophage reporter, Hulsmans *et al*.^10^ reported a striking abundance of macrophages along the mouse AV conduction axis and proposed that resident macrophages modulate AV conduction by forming spatially restricted Cx43-containing gap junctions with myocytes in the distal AV node. In that study, genetic manipulation or ablation of CX3CR1-expressing cells impaired AV node function, leading to the widely cited conclusion that macrophages are required for normal AV conduction.^10^ The work rapidly became a flagship example of “cardio-immunology” in action, catalysing broad interest in immune cells as active, dynamic regulators of cardiac excitability and, by extension, shaping subsequent experimental priorities and funding decisions across the field.^11,12^ Three considerations motivated a re-examination of this phenomenon:(i)First, transcriptomic datasets indicate that cardiac macrophages express inward-current channel genes at low levels,^10,13,14^ consistent with patch-clamp evidence that they lack inward currents^15^ indicating that any direct electrical influence would be electrotonic. **(ii)** Because macrophages are relatively depolarised (resting membrane potentials reported in the range ∼−55 to −20 mV^15^), basic biophysics predicts that coupling to an automatic nodal myocyte (maximum diastolic potential near ∼−60 mV^16^) would shift myocyte diastolic potential positively, reduce the the driving force for key voltage-dependent inward currents and thereby suppress excitability and conduction. **(iii)** The AV node is generally considered Cx43 poor. We therefore addressed the proposed mechanism using three complementary approaches. First, we implemented contemporary biophysical models of mouse AV node myocytes and macrophage subtypes to predict the consequences of coupling AV node pacemaking and action potential conduction. Second, we mapped Cx43 relative to HCN4-expressing AV node myocytes and CD68-expressing macrophages in mouse AV junction using high-resolution immunohistochemistry and optical tissue clearing, and we performed analogous analysis in non-failing and failing human penetrating bundle (a region of the human AV junction in which Cx43 expression has been reported^17^). Third, we tested functional necessity by depleting resident macrophages in healthy mice using CSF1R inhibition and quantifying AV conduction indices *in vivo* and *ex vivo*.

## METHODS

Detailed experimental protocols, including human tissue acquisition and processing, animal models and interventions, computational simulations, histology, immunolabelling, imaging and electrophysiological recordings, are provided in the Supplemental Information. The full dataset is available from the corresponding authors on reasonable request.

## RESULTS

### Macrophage coupling disrupts AV node excitability, automaticity and conduction *in silico*

Hulsmans *et al*.^10^ provided simulations in support of the view that macrophages facilitate AV node conduction in mouse hearts, but the only models available to them at the time were an early model of the rabbit AV node myocyte^18^ based on disparate data sources, and a model of a fibroblast rather than a macrophage.^19^ We assessed the plausibility of their mechanism using contemporary *in silico* models. AV node myocytes were simulated with the Bartolucci HCN4+ mouse AVN model,^20^ and macrophages with subtype-specific mouse cardiac macrophage models of Simon-Chica *et al*.^15^ Shown here are results from the Type 0 macrophage model and results from the other macrophage phenotype models are provided in Supplemental Figs. S1 and S2. We first simulated 1:1 AV node myocyte-macrophage coupling during physiologically relevant pacing (350–550 beats/min), increasing gap-junctional conductance (G_gap_) from 0 to 7 nS (Fig. 1A-D, 550 beats/min shown). In these simulations, increasing G_gap_ resulted in a prolongation of the myocyte action potential (action potential duration at 90% repolarization, APD_90_, increased by ∼13 ms at G_gap_ of 3 nS; Fig. 1A,B) as well as a depolarizing shift in the diastolic potential (Fig. 1A,C). The prolongation of the myocyte action potential is largely the result of inward gap junctional current, *I*_GapMΦ_ (dashed grey line in Fig.1D), during the second half of the action potential when the myocyte is repolarizing and the myocyte membrane potential becomes more negative than the macrophage membrane potential (*I*_GapMΦ_ is outward during the overshoot of the myocyte action potential in the first half when the myocyte membrane potential is more positive than the macrophage membrane potential).

**Figure 1.**
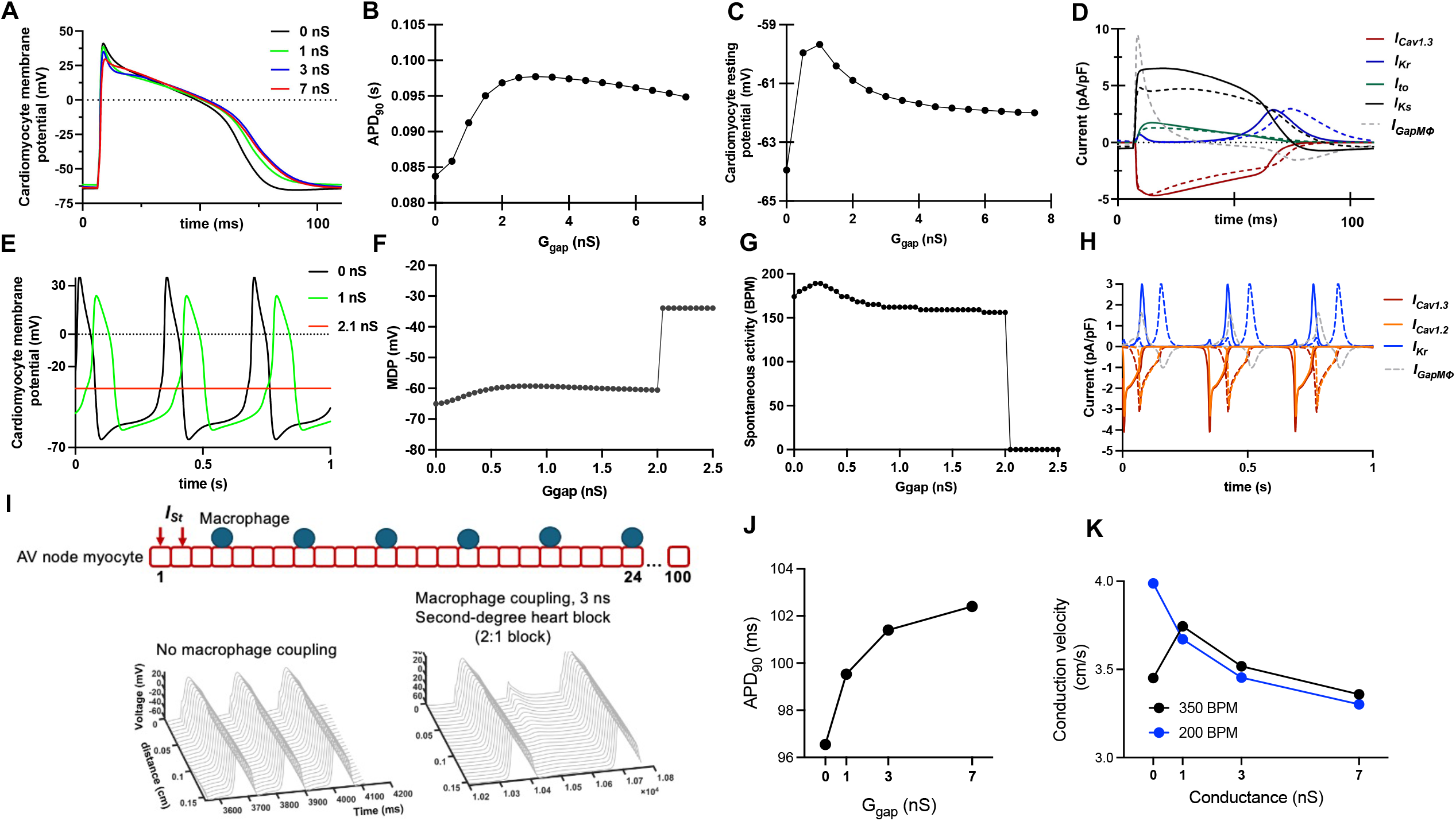
Computational modelling predicts macrophage coupling suppresses AV node excitability, automaticity and conduction. (A)Paced AV node myocyte action potentials (550 beats/min) with increasing myocyte–macrophage coupling conductance (Ggap) to a Type 0 macrophage (A–K). (B) Relationship between myocyte APD90 and Ggap (550 beats/min). (C) Relationship between myocyte resting/diastolic potential and Ggap.(D) Ionic currents underlying paced activity (*I*_Ks_, *I*_Cav1.3_, *I*_Kr_, *I*_to_ and *I*_GapMϕ_). Solid lines, no coupling (0 nS); dashed lines, coupling (3 nS). (E) Representative spontaneous AV node myocyte activity during a ramp increase in Ggap (0–7.5 nS over 20 s). (F) Maximum diastolic potential (MDP) as a function of Ggap during the ramp protocol. (G) Spontaneous firing rate as a function of Ggap during the ramp protocol. (H)Ionic currents underlying spontaneous activity during coupling (*I*_Cav1.2_, *I*_Cav1.3_, *I*_Kr_ and *I*_GapMϕ_). Solid lines, no coupling; dashed lines, coupling (conductance as indicated).(I)One-dimensional strand model schematic (24 or 100 AV node myocytes) with one macrophage coupled in parallel to every fourth myocyte; the first two myocytes are paced (*I*_st_). Space–time plots show stable second-degree (2:1) block with coupling (3 nS) versus no coupling (0 nS) in a 24-cell strand. (J) Relationship between APD90 and Ggap in a 100-cell strand (350 beats/min).(K) Relationship between conduction velocity and Ggap in a 100-cell strand at 350 and 200 beats/min.

Automaticity is a defining feature of AV node myocytes. To test how coupling affects spontaneous pacemaking in AV node myocytes, we ramped G_gap_ from 0 to 7.5 nS over 20 s (Fig. 1E-H). As G_gap_ was increased the maximum diastolic potential of the myocyte decreased from −65 to −62 mV (Fig. 1E,F), whereas pacemaking initially accelerated from 174 to 189 beats/min at 0.25 nS and then slowed to 168 beats/min at 0.6 nS (Fig. 1E,G). Above G_gap_ of 2.0 nS, pacemaking failed and the membrane potential settled at −34 mV (Fig. 1E-G). Notably, at G_gap_ of 3 nS the AV node myocyte was quiescent (Fig. 1E-G); 3 nS was the G_gap_ assumed by Hulsmans *et al*.^10^ at a macrophage-myocyte junction (assuming three Cx43 contact points between a macrophage and a myocyte and a G_gap_ of ∼1 nS per contact point). The depolarization of the diastolic membrane can be attributed to the inward *I*_GapMΦ_ discussed and the slowing of pacemaking can be attributed to a progressive loss of *I*_Cav1.3_ as a result of inactivation caused by the depolarization (Fig. 1H).

The AV node’s primary role is action potential conduction and therefore we next tested how macrophage coupling influences conduction in a one-dimensional (1D) strand model of the AV node. We used 1D strands of 24 or 100 AV node myocytes, with one Type 0 macrophages coupled in parallel to every fourth myocyte; the first two myocytes were paced until a steady-state was reached (Fig. 1I, top panel). In the absence of coupling, the conduction velocity was ∼3.5 cm/s (like that determined experimentally^9^). G_gap_ was then set to 3 nS (assuming three Cx43-mediated myocyte-to-macrophage contact points). With G_gap_ of 3 nS and a stimulation rate of 350 beats/min, second-degree (2:1) block developed (Fig. 1I, lower panels) and persisted for tens of seconds (Supplemental Fig. S2E). Macrophage-myocyte coupling prolonged APD_90_ (Fig. 1J) as in the previous simulations; this will increase the AV node refractory period and this can explain the 2:1 block at a stimulation rate of 350 beats/min. It is well known that the conduction velocity of the AV node is rate-dependent^21^ aand, because of the effective halving of the conducted rate during 2:1 block at a stimulation rate of 350 beats/min, the conduction velocity initially increased. However, at a stimulation rate of 350 beats/min, with further increases in G_gap_ the conduction velocity monotonically declined (Fig. 1K).

With a lower stimulation rate of 200 beats/min, there was 1:1 conduction throughout (the increase in APD_90_ had no impact because of the longer diastolic interval) and only the monotonic decline in the conduction velocity was observed (Fig. 1K). Together, these modelling analyses predict that macrophage–myocyte electrical coupling would suppress AV nodal automaticity and conduction rather than facilitate conduction.

### No evidence of Cx43-mediated macrophage–myocyte coupling in the mouse AV conduction axis

Because the coupling configuration proposed to facilitate conduction^10^ produced the opposite effect in our simulations, we next determined how frequently Cx43 contact points between macrophages and nodal myocytes can be detected in the mouse AV conduction axis. Although CX3CR1 reporters are widely used to label tissue-resident macrophages^10,22,23^ on closer inspection endogenous CX3CR1 expression is detectable in HCN4-expressing AV node myocytes (Fig. 2A,B) that exhibit *I*_f_ (Fig. 2C), indicating that CX3CR1 is not restricted to macrophages in this tissue (Figs. 2B and S3A-B). Consistent with this, reanalysis of published single nucleus RNAseq data of the mouse sinoatrial node^13^ demonstrates that the transcript expression of *Cx3cr1* is higher in nodal myocytes than in macrophages (Fig. S2D), and protein expression of CX3CR1 has been previously reported in myocytes of the rodent and human working myocardium at baseline.^24,25^ The limited cellular specificity of the CX3CR1 reporter in the AV junction is also apparent in previous studies in which the CX3CR1 signal is prominent in the central fibrous body (Fig. 1A in Hulsmans *et al*.^10^) – a collagenous structure not generally regarded as being densely populated by macrophages. Given an apparent lack of macrophage specificity of CX3CR1 in the mouse AV node, we used CD68 to identify macrophages.^26,27^ CD68 is a lysosomal glycoprotein highly enriched in macrophage lineage cells, is robustly preserved in fixed cardiac tissue, and was not expressed in in HCN4-positive AV node myocytes in our material (Fig. S3A). To assess the number of Cx43 contact points per AV node myocyte *in situ*, 10 μm serial tissue sections were cut through preparations encompassing the mouse AV conduction axis (Fig. 2A), extending from the inferior nodal extension (INE) through the compact AV node (CN) and penetrating bundle (PB) to the proximal bundle of His (BoH) and immunolabelled for HCN4 and Cx43 (Fig. 2D, extended seriesis shown in in Fig. S3). Across this axis, Cx43 signal was absent from HCN4-positive myocytes in the INE, CN, PB and proximal BoH, and was confined to surrounding HCN4-negative myocardium (Fig. 2D). To directly assess proximity between macrophages and Cx43 within nodal tissue, the entire AV conduction axis was triple-labelled for HCN4, Cx43 and CD68. Representative high-power images of the distal penetrating bundle and compact node are shown in Fig. 2E–F. Manders overlap coefficients quantified co-localisation between Cx43 and HCN4, and between Cx43 and CD68, within defined components of the HCN4-positive conduction axis; in both cases overlap was low (Fig. 2G). Finally, to exclude the possibility that the absence of signal overlap reflected sectioning artefact, we performed optical clearing and whole-mount immunolabelling of the mouse AV conduction axis for HCN4, Cx43 and CD68 (Fig. 2H). Cleared-tissue imaging recapitulated the serial-section findings: the HCN4-positive AV conduction axis lacked detectable Cx43 signal, supporting the conclusion that a Cx43-based macrophage– myocyte coupling substrate is not evident in the mouse AV node under basal conditions.

**Figure 2.**
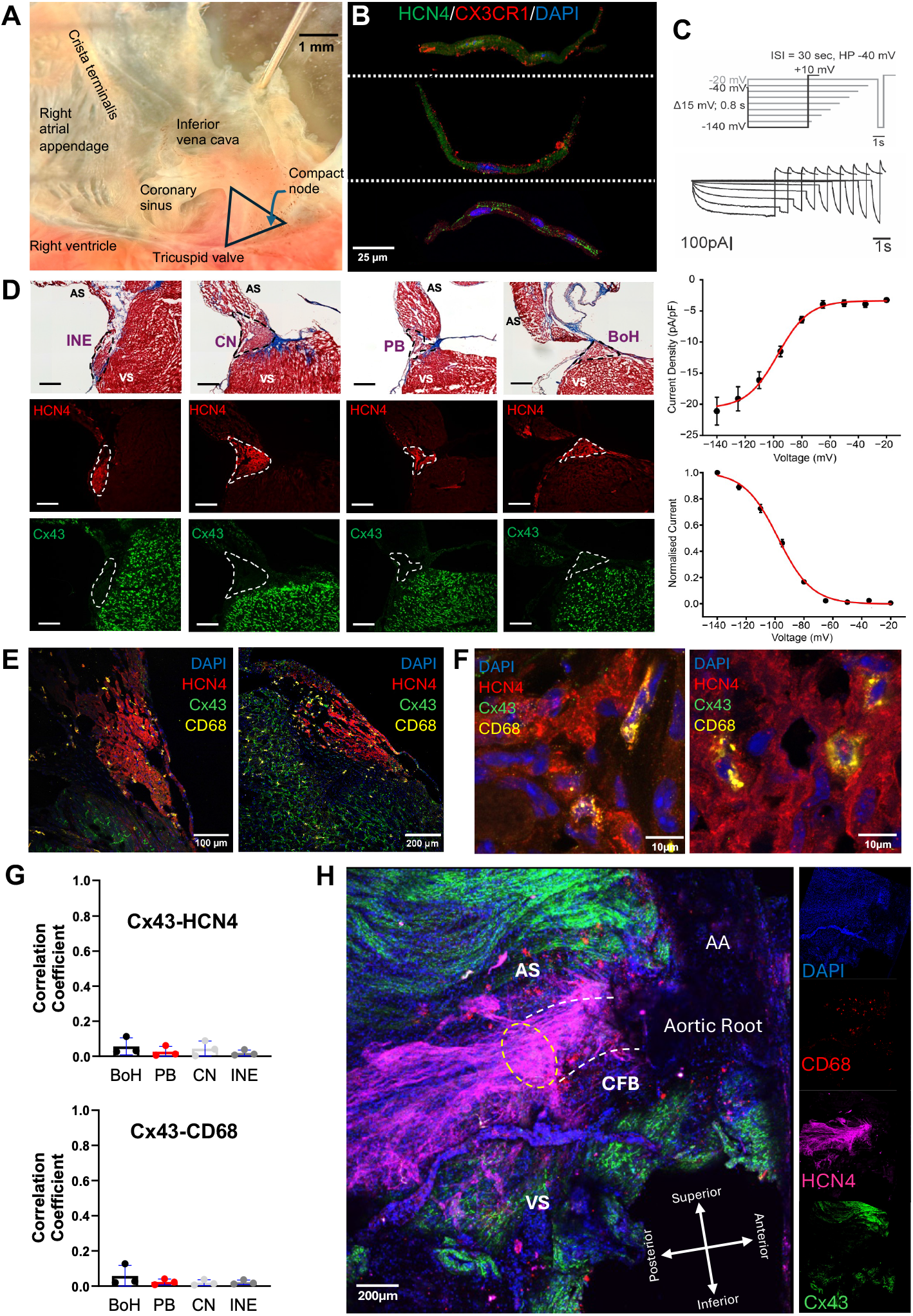
HCN4-positive mouse AV conduction tissue lacks Cx43 and shows minimal macrophage–Cx43 overlap. (A) Right atrial endocardial preparation indicating the Triangle of Koch region analysed. (B) Isolated AV node myocytes immunolabelled for HCN4, CX3CR1 and DAPI (nuclei). (C) *I*_f_ recordings from AV node myocytes showing the voltage-clamp protocol, representative traces, mean current density–voltage relationship and activation curve (n=16 cells from 4 mice; mean ± SD; Boltzmann fit shown). (D) Serial long-axis sections through the AV conduction axis: Masson’s trichrome (top) and matched immunolabelling for HCN4 and Cx43 (bottom); extended series in Fig. S3. Scale bars, 200 μm. (E,F) High-power images of penetrating bundle (E) and compact node (F) immunolabelled for HCN4, Cx43, CD68 and DAPI. (G) Manders overlap coefficients for Cx43 with HCN4 (top) and with CD68 (bottom) across AV conduction components (3 biological replicates; mean ± SD). (H) Optically cleared AV conduction axis immunolabelled for HCN4, Cx43 and CD68 (DAPI). AA, ascending aorta; AS, atrial septum; BoH, bundle of His; CFB, central fibrous body; CN, compact node; INE, inferior nodal extension; PB, penetrating bundle; VS, ventricular septum.

### Cx43-mediated macrophage–myocyte coupling is not discernible in the human distal conduction system

We next assessed whether macrophages in the human AV conduction system are equipped to participate in Cx43 coupling. Interrogation of published human AV node multi-omic datasets (single-cell/single-nucleus RNA-seq with spatial transcriptomics) indicates that AV node macrophages exhibit near-zero expression of GJA1 (Cx43 transcript), in contrast to abundant GJA1 expression in working myocardium-derived myocytes.^14^ This dissociation suggests that macrophages of the human AV conduction system are transcriptionally unlikely to form Cx43 gap junctions. To complement this with direct protein-level assessment, we examined the human penetrating bundle (Fig. 3), a region previously reported to express Cx43.^17^ Tissue from non-failing (n=3) and failing (n=3) human hearts was analysed (Fig. 3A-B). As expected, Cx43 puncta were abundant and CD68-expressing macrophages were present within and around the penetrating bundle in both sets of hearts (Fig. 3C). The tissue was systematically screened by high-resolution confocal microscopy; instances in which Cx43 puncta lay in close apposition to CD68-expressing cells were documented (examples from non-failing and failing hearts are shown in the lower panels of Fig. 3C), but such events were exceedingly rare and true signal colocalisation was not observed. Correlation coefficients were low in both groups and remained near zero; failing hearts showed a slight numerical increase but no meaningful colocalisation (Fig. 3D). Taken together, these findings indicate that Cx43-mediated electrical coupling between macrophages and AV node myocytes is unlikely in the human AV conduction axis.

**Figure 3.**
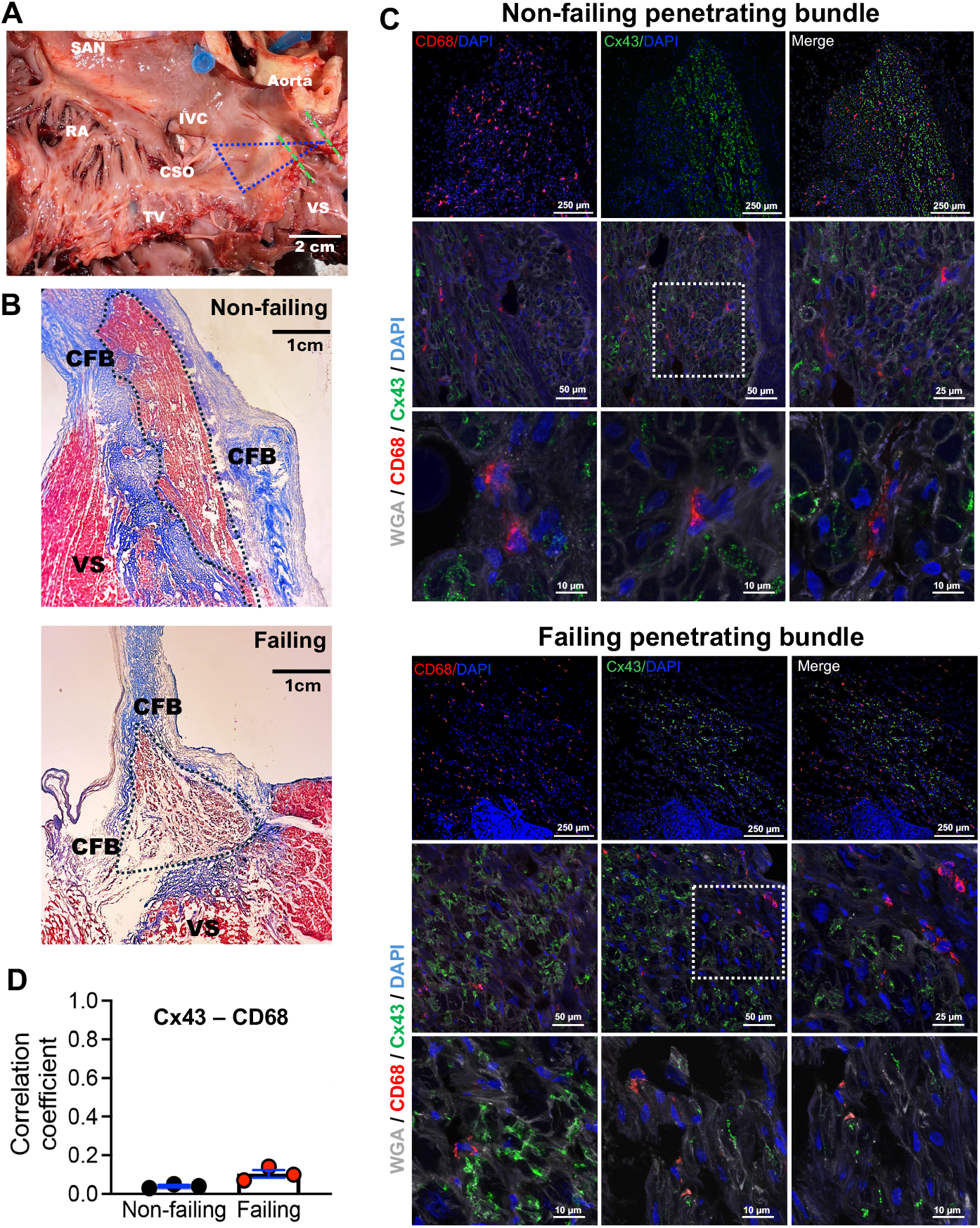
Cx43 does not colocalise with CD68 in the human penetrating bundle. (A) Human right atrial endocardial surface indicating the Triangle of Koch and Cx43-positive regions of the penetrating bundle identified by serial immunolabelling. (B) Penetrating bundle sections from non-failing and failing hearts identified by serial sectioning and Masson’s trichrome staining. (C) Immunolabelling for Cx43 and CD68 with WGA (cell boundaries) and DAPI (nuclei) in non-failing (top) and failing (bottom) penetrating bundle, shown at low and high magnification with regions of interest indicated. (D) Manders overlap coefficients for Cx43 and CD68 in non-failing and failing penetrating bundle (n=3 hearts per group; mean ± SD). Abbreviations: CFB, central fibrous body; CSO, coronary sinus ostium; IVC, inferior vena cava; RA, right atrium; SAN, sinoatrial node; TV, tricuspid valve; VS, ventricular septum.

### Macrophage depletion does not impact AV node conduction

Direct coupling notwithstanding, are macrophages required for AV node conduction? To test whether resident cardiac macrophages are required to maintain AV node conduction under normal conditions, we depleted macrophages pharmacologically using PLX5622, a small-molecule inhibitor of colony-stimulating factor 1 receptor (CSF1R) that depletes macrophage populations by blocking CSF1R-dependent survival signalling.^28^ Adult mice were fed a PLX5622-formulated diet (1200 ppm) for 21 days, after which AV node function was assessed *in vivo* by surface ECG recording and *ex vivo* by electrophysiological study of Langendorff-perfused hearts (Fig. 4). Macrophage depletion was confirmed by flow cytometry^29^ of F4/80^+^cardiac macrophages isolated from micro-dissected Triangle of Koch tissue, demonstrating a 96.4% reduction in PLX5622-treated animals compared with controls (Fig. 4A–B). Depletion within the AV junction was further supported by immunolabelling of cryosections for HCN4 and CD68, which showed marked loss of CD68-expressing cells in PLX5622-treated preparations relative to controls (Fig. 4C). *In vivo*, PLX5622 treatment did not alter ECG intervals (Fig. 4D). Despite near-total macrophage depletion, the PR interval was unchanged (35.7±0.77 ms and 35.6±0.81 ms in treated and control mice, respectively), and no second- or third-degree AV block was observed (Fig. 4D). Heart rate, QRS duration and QTc interval were also comparable between groups (Fig. 4D). We next assessed AV nodal conduction *ex vivo* using Langendorff perfusion and programmed stimulation (Fig. 4E). Conduction remained robust in macrophage-depleted hearts: Wenckebach cycle length (WBCL) was unchanged between PLX5622-treated and control groups, indicating preserved AV nodal conduction at high atrial pacing rates (Fig. 4E). Conduction remained robust in macrophage-depleted hearts: the PR interval measured at a fixed pacing cycle length (120 ms, allowing assessment of the PR interval independent of sinus rate), the Wenckebach cycle length, and the AV node effective refractory period were all unchanged between the PLX5622-treated and control groups, indicating preserved AV node conduction (Fig. 4E). These data support the hypothesis that resident macrophages are not required for maintaining normal AV conduction.

**Figure 4.**
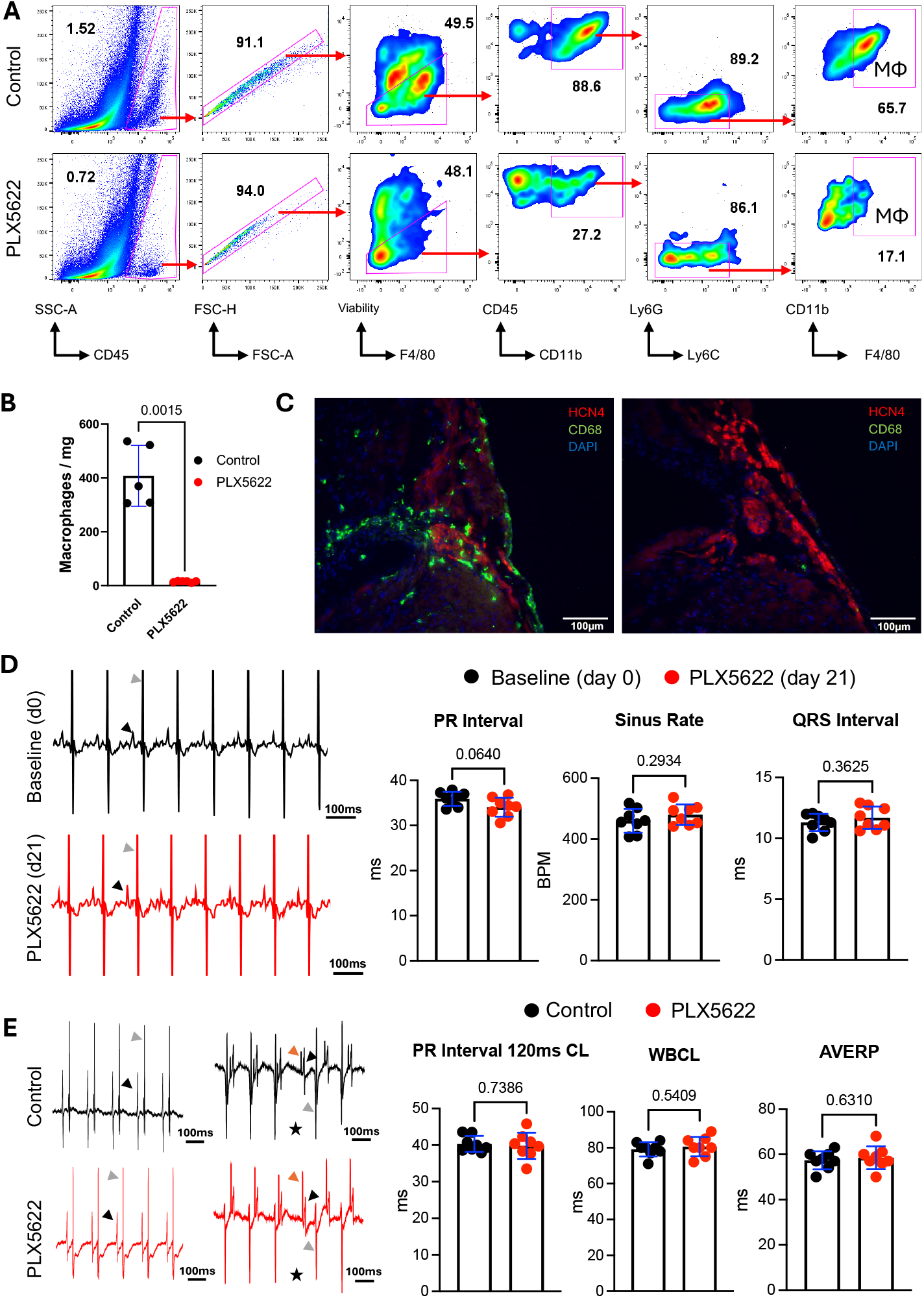
CSF1R inhibition depletes cardiac macrophages without altering AV node conduction. (A) Representative flow cytometry gating for macrophage quantification in micro-dissected Triangle of Koch tissue from control mice or mice fed a PLX5622-formulated diet (1200 ppm). (B) Macrophages per mg Triangle of Koch tissue quantified by flow cytometry (n=5 per group; mean ± SD with individual points; Welch’s t test; 96.4% reduction). (C) Penetrating bundle sections from control and PLX5622-treated mice immunolabelled for HCN4, CD68 and DAPI. (D) *In vivo* ECG example traces and group data for PR interval, heart rate and QRS duration at baseline and after 21 days of PLX5622 diet (n=8; mean ± SD with individual points; Welch’s t test). (E) *Ex vivo* pseudo-ECG example traces and PR interval at paced CL 120 ms, Wenckebach cycle length (WBCL) and AV node effective refractory period (AVERP) in Langendorff-perfused hearts from control and PLX5622-treated mice. Black arrowheads indicate P waves, grey arrowheads indicate QRS complexes and black stars indicate dropped QRS complexes. (n=8; mean ± SD with individual points; Welch’s t test).

## DISCUSSION

The proposal that resident cardiac macrophages facilitate atrioventricular (AV) node conduction via Cx43-containing gap junctions offered an eye-catching bridge between cardiac electrophysiology and immunology. However, the combined biophysical, anatomical and functional evidence presented here places constraints on a Cx43 coupling mechanism as an explanation for basal AV node function.

Across experimental systems, heterocellular coupling between excitable cardiomyocytes and non-excitable cells is most consistently associated with electrotonic loading, depolarisation of the myocyte and conduction slowing, rather than improved excitability.^30-32^ Mechanistically, coupling a relatively depolarised macrophage to an AV node myocyte imposes an electrotonic load that shifts the myocyte to more positive potentials. Because AV node myocytes are small (capacitance ∼20 pF^16^, comparable to a macrophage’s ∼18 pF^15^) and macrophages lack fast inward currents, the coupled macrophage behaves as a current sink; as shown by Fig. 1 even modest coupling can slow pacemaker firing and, above a threshold, terminate spontaneous activity. In earlier modelling,^10^ inference was largely based on resting potential and APD, whereas AV node–specific functional outputs (pacemaking, propagation, WBCL/AVERP) are needed to evaluate facilitation versus suppression.

Our immunolabelling data also limit the plausibility of a direct Cx43 coupling mechanism within the AV node. The AV conduction axis is characterised by low-conductance connexins, and Cx43 is absent in the HCN4-defined nodal myocardium in mice (Fig. 2). Against that background, rare proximity between macrophages and Cx43 puncta (undetectable in our study) is not, on its own, evidence for a physiologically meaningful coupling architecture. Our structural analyses suggest that such a substrate is not a prominent feature of the AV node under basal conditions, and that macrophage–Cx43 co-localisation events are negligible-to-absent even in the human penetrating bundle where Cx43 is abundant. Prior studies^10,33^ have examined AV node conduction after macrophage depletion using a range of genetic and pharmacologic tools, with divergent results.

Germline models that lack macrophages from development – e.g. Csf1^op/op^ mice – show first-degree AV block at baseline, raising the possibility that developmental remodeling or broader structural abnormalities contribute to the phenotype, rather than acute loss of macrophage–myocyte coupling *per se*. Broad diphtheria-toxin ablation of CD11b^+^myeloid cells can precipitate complete heart block,^10^ but DTR-based ablation can provoke marked inflammatory activation after massive cell death, complicating attribution of conduction phenotypes to macrophage loss per se.^34^ More recently, CSF1R-targeted approaches have provided a complementary view. In LysM^Cre^ × Csf1r^LsL-DTR^ (MM^DTR^) mice, diphtheria-toxin–induced depletion of CSF1R-expressing myeloid cells substantially reduced cardiac macrophages yet leaves baseline ECG intervals, including PR duration, unchanged.^33^ In this context, preserved AV nodal function after CSF1R inhibition is most consistent with macrophages not being required to maintain basal conduction in healthy adult animals. A further interpretive constraint is the cellular specificity of tools used to identify and manipulate macrophages in the AV junction. CX3CR1 is expressed by HCN4-expressing AV node myocytes (Fig. 2B), and single-nucleus RNAseq datasets detect *Cx3cr1* transcripts across multiple cardiac cell types with high expression in nodal myocytes (Supplementary Fig. S3D). Accordingly, CX3CR1-driven optogenetic stimulation or gene deletion cannot be assumed to be macrophage-specific in the AV junction without additional targeting controls.

Based on the information given here, direct macrophage–myocyte Cx43 gap-junction coupling is unlikely to be a basal mechanism sustaining AV nodal conduction. This does not diminish the broader importance of macrophages in conduction system biology; rather, it shifts emphasis toward context-dependent, indirect mechanisms. In disease or stress states, macrophages can modulate conduction through paracrine signalling that influences myocyte–myocyte coupling, tissue structure and ion channel function. Amphiregulin-dependent preservation of cardiomyocyte Cx43 phosphorylation under acute stress provides one example,^35^ and macrophage-derived mediators implicated in heart failure remodelling (including galectin-3) provide another.^36,37^ Under pro-inflammatory conditions (e.g., LPS exposure, obesity and infection), macrophage Cx43 can be upregulated but reported effects are predominantly mediated via hemichannels (such as ATP release) rather than formation of intercellular gap junctions.^38-42^ AV nodal macrophages may therefore be best viewed as contextual regulators: quiescent with respect to direct electrical coupling at baseline, but capable of influencing conduction indirectly through inflammation-responsive signalling pathways in pathology. Future work should define the conditions under which these interactions become maladaptive and whether macrophage-derived pathways offer tractable therapeutic targets for conduction system disease.

## ACKNOWLEDGEMENTS

This work was funded by a British Heart Foundation (BHF) Intermediate Basic Science Research Fellowship (FS/19/1/34035; FS/EXT/24/35026) BHF International Cardiovascular Research Partnership Award (IA/F/24/275114), BHF project grants (PG/22/10919; PG/25/12197) and a BHF Clinical Research Training Fellowship (FS/CRTF/23/24469) to AD. ADS, MRB and MEM received funding from *a Fondation Leducq* award (TNE FANTASY 19CV03).

## Notes

### Competing Interest Statement

The authors have declared no competing interest.

