## Supplement for "Resident cardiac macrophages are not required for normal atrioventricular node conduction"

### Data Supplement

### Methods

#### Animals

Work was conducted in accordance with The UK Animals (Scientific Procedures) Act (1986) under project licence PP3219770. C57BL/6J mice aged 10-12 weeks were used. $Cx_3cr1^{GFP/+}$ mice[1] were provided by Professor Matthew Hepworth at the University of Manchester. Mice were housed at the Imperial College London CBS Facility or the University of Manchester Biological Services Facility, providing temperature-controlled housing, 12:12 hour light-dark cycle, regular monitoring and free access to food, water and environmental enrichment items in both locations.

#### Human subjects

Human cardiac tissue was obtained from the Imperial College Human Tissue Programme (HTP), approved under the UK Human Tissue Authority (HTA) licence: donor (non-failing) hearts were provided under IRAS project ID 189069 (REC reference: 16/LO/1568) and failing samples collected under IRAS project ID 264059 (REC reference: 19/SC/0257) from the Royal Brompton and Harefield Hospitals Cardiovascular Biobank. All samples were collected and used in accordance with the Human Tissue Act (2004) and institutional ethical approval. Written informed consent was obtained from all donors (or their next of kin, where applicable).

| **Failing (*n* = 3)** | | | | | | | |
|---|---|---|---|---|---|---|---|
| **ID** | **Age** | **Sex** | **HF Aetiology** | **CVD** | **Hypertension** | **DM** | **Smoking** |
| F1 | 46 | M | DCM | Y | N | N | Y |
| F2 | 68 | M | DCM | Y | Y | N | N |
| F3 | 60 | F | Sarcoidosis | N | N | Y | N |
| **Non-failing (*n* = 3)** | | | | | | | |
| **ID** | **Age** | **Sex** | **Cause of Death** | **CVD** | **Hypertension** | **DM** | **Smoking** |
| NF1 | 71 | F | ICH | N | N | N | N |
| NF2 | 70 | F | ICH | N | N | N | N |
| NF3 | 76 | M | ICH | N | Y | N | Y |

**Table S1. Donor demographic data.** Key cardiovascular and lifestyle risk factors are presented. CVD - cardiovascular disease; DM - diabetes mellitus; ICH - intracranial haemorrhage; HF – heart failure.

#### Numerical modelling of AV myocyte automaticity and conduction

We employed the recently developed models of AV myocytes by Bartolucci et al[2] and cardiac macrophages subtype models by Simon-Chica *et al.*[3] The Bartolucci model accurately reproduces evoked as well as spontaneous action potentials of HCN4-expressing AV node myocytes and incorporates key ionic currents including the HCN-mediated funny current ($I_f$), the $Ca_v1.2$-mediated ($I_{Cav1.2}$) and $Ca_v1.3$-mediated ($I_{Cav1.3}$) L-type $Ca^{2+}$ currents, the $Ca_v3.1$-mediated T-type $Ca^{2+}$ current ($I_{Ca,T}$), the cardiac $Na_v1.5$-mediated TTX-insensitive $Na^+$ current ($I_{Na}$), as well as a TTX-sensitive $Na^+$ current and inward-rectifier and delayed outward rectifier $K^+$ currents. The Simon-Chica models emulate four distinct CX3CR1-expressing cardiac macrophage electrophysiological phenotypes (Types 0–2 and Type 1+2) based on their expression of outward $K_v1.3$, $K_v1.5$ and $K_v2.1$ $K^+$ currents. All macrophage models include Cx43-mediated junctional coupling. To simulate coupling between AV myocytes and macrophages *in silico*, we first integrated the general model of single AV myocyte with models of subtypes of macrophages (Type 0-2 and 1+2) as reported in Simon-Chica *et al,*[3] to create a myocyte-macrophage 1:1 coupling cellular model. Differential equations, kinetic parameters and membrane conductances employed to simulate AV myocyte and macrophages were the same as in Bartolucci *et al.* and Simon-Chica *et al.*, respectively.

Coupling AV myocyte (Myo) and macrophage (Mϕ) in the 1:1 cellular coupling model was simulated assuming linear resistive coupling. Myo-Mϕ intercellular connection gap current ($I_{Gap}$) was calculated according to the equations:

$$IGapMyo = (\frac{GgapM\phi}{CmMyo})(VMyo - VM\phi)$$

$$IGapM\phi = (\frac{GgapM\phi}{CmM\phi})(VM\phi - VMyo)$$

where $G_{gap}$ is the gap junction conductance, $V_{Myo}$ and $V_{M\phi}$ are the myocyte and macrophage membrane voltage, respectively and assuming that $G_{gapM\phi} = G_{gapMyo}$
The membrane potential of connected myocyte and macrophage was calculated according to the equation:

$$Cm\frac{dVMyo}{dt} = -IionMyo - IGapMyo$$

for myocyte membrane potential and:

$$Cm\frac{dVM\phi}{dt} = -IionM\phi - IGapM\phi$$

for macrophage membrane potential, where $C_m$ is the cell capacitance, V is the membrane voltage, $I_{ion}$ is the sum of endogenous ionic currents of myocyte and macrophage and $I_{Gap}$ is the current flowing through gap junction. The myocyte membrane potential under pacing stimulation was calculated according to the equation:

$$Cm\frac{dVMyo}{dt} = -IionMyo - IGapMyo + Ist(t)$$

where $I_{st}$ is the current stimulation. To predict the effect of macrophages on myocyte action potential at 90% repolarisation ($APD_{90}$) in the 1:1 model, the myocyte model was paced at 450 and 550 beats per minute with a pulse stimulus of 600 pA amplitude and 1 ms duration as in Bartolucci et al[2]. Simulations were run for 20 s to reach equilibrium.

To predict the effects of coupling between AV myocytes and macrophages, we built a monodimensional (1D) strand of n1 to n24 or n1 to n100 AV node myocytes (see Fig. 2A). The time course of myocytes action potential was calculated by the general partial differential equation:

$$Cm\frac{dV(x,t)}{dt} = -Iion + Icouple + Ist(x,t)$$

Where x is the space, t is the time, $I_{ion}$ is the sum of ionic currents of the given n myocyte and $I_{couple}$ is the coupling current between myocytes. To measure conduction velocity along the 1D strand model, the first two myocytes (n1-n2) were simultaneously paced at 200 or 350 beats per minute with a pulse stimulus of 200 pA amplitude and 5 ms duration. Differential equations were solved using the numerical differentiation method as in Bartolucci *et al.*[2] with a maximum time step of 1 ms. Partial differential equations were integrated using the Method of Lines (MOL).[4] Simulations and integrations were run using Matlab (ver. 2025B, Mathworks Inc.). Simulation results were analysed using custom made Matlab script automatically detecting and reporting frequency of action potentials and the following action potential parameters: AP threshold, $APD_{90}$, maximum diastolic potential MDP, the maximum upstroke potential (MUP). Conduction velocity of action potential along the strand was calculated by determining the distance (in cm) and time delay (in seconds) between the MUP of myocyte n5 and myocyte n10 (See Fig. 2A). Simulation results were plotted using Prism (ver. 10, GraphPad), or Matlab. The model code is available for download at Github public server with link: https://github.com/CARDIAC-RHYTHM-PHYSIOPATHOLOGY/Myocyte-Macrophage-Coupling. Results and code were independently verified by coauthors at the University of Leeds.

**Isolation of mouse AV node myocytes**

Isolation and electrophysiological study of mouse AV node myocytes was performed as described previously.[5] Eight to twelve-week-old mice were euthanized by cervical dislocation. Beating hearts were quickly dissected and transferred to pre-warmed Tyrode's solution (140 mM NaCl, 5.4 mM KCl, 1 mM $MgCl_2$, 1.8 mM $CaCl_2$, 5 mM HEPES-NaOH (pH 7.4), and 5.5 mM D-glucose in a silicone-lined Petri dish. The AV node region was isolated and transferred to pre-warmed (37°C) Tyrode's "low $Ca^{2+}$" solution containing 140 mM NaCl, 5.4 mM KCl, 0.5 mM $MgCl_2$, 0.2 mM $CaCl_2$, 1.2 mM $KH_2PO_4$, 50 mM taurine, 5 mM HEPES-NaOH (pH 6.9), and 5.5 mM D-glucose) in a 2 ml Eppendorf vial. Bovine serum albumin (BSA, 1 mg ml−1, Merck KGaA, Germany), elastase (18.4 U ml−1, Merck KGaA, Germany), collagenase B (0.3 U ml−1, Roche Diagnostics, Germany), and protease (1.8 U ml−1, Merck KGaA, Germany) were added to the Eppendorf vial and incubated at 37°C in the heat block for 26 – 28 mins at 36 °C. The AV node tissue contained in the vial was then centrifuged at 200 xg for 2 mins at 4°C and the supernatant was removed. The tissue was subsequently washed twice with Tyrode's low $Ca^{2+}$ solution, and then twice with ´Kraftbrühe´ (KB) buffer containing: 80 mM L-glutamic acid, 25 mM KCl, 3 mM MgCl2, 10 mM KH2PO4, 20 mM taurine, 10 mM HEPES-KOH (pH = 7.4), 0.5 mM EGTA, and 10 mM D-glucose. Finally, the tissue was resuspended in 350 µl KB buffer and set to recover for 3-4 hrs at 4°C. After recovery, the AV node tissue was acclimated to room temperature for 15 min before being mechanically dissociated by pipetting 4 to 8 times with a flame forged Pasteur's pipette. 50 µl of cell suspension was added to poly-l-lysine (PLL)-coated coverslips and allowed to set for 15 mins to ensure proper cell attachment to PLL substrate.

**Electrophysiological recording of $I_f$ in AV node myocytes**

For patch-clamp recording of the funny current ($I_f$), AVN cells were set to the bottom of PLL-coated coverslip. $Ca^{2+}$ was then reintroduced into the KB solution to restore the physiological extracellular $Ca^{2+}$ concentration. Cells were perfused with Tyrode's solution at 32 °C. Patch-clamp pipettes were pulled to reach final resistance of 3-5 MΩ when filled with an intracellular solution containing: 90 mM potassium aspartate, 10 mM NaCl, 2.0 mM $MgCl_2$, 2.0 mM $CaCl_2$, 5.0 mM EGTA, 2.0 mM $Na_2$-ATP, 0.1 mM $Na_2$-GTP, and 5.0 mM creatine phosphate (pH 7.2). During the voltage clamp recording, the cells was perfused with modified Tyrode's solution containing $BaCl_2$: 140 mM NaCl, 5.4 mM KCl, 1 mM $MgCl_2$, 1 mM $CaCl_2$, 5 mM HEPES, 5 mM D-glucose (pH=7.4). 2 mM $BaCl_2$ and 0.3 mM $CdCl_2$, were added to inhibit inward rectifier $K^+$ current ($I_{kir}$) and $Ca^{2+}$ channels, respectively. Recordings were performed at 32 °C. $I_f$ was recorded using the whole-cell variation of patch-clamp technique by applying hyperpolarizing voltage steps of variable duration, to steady-state current activation, with a protocol consisting of from -140mV to -50 mV in -15mV increments every 4.5 s from a holding potential of -40mV, before applying a test potential of -140mV to current full activation for 500ms. Data were recorded using Clampex 11 (Molecular Devices). The $I_f$ activation curve was calculated by plotting the normalized steady-state current amplitude versus the applied voltage step after automatic linear leakage subtraction. The curve was then fitted to the Boltzmann equation. $I_f$ isochronal current-to-voltage relationship was obtained at different hyperpolarized voltages from -140mV to -80mV. Current density was obtained by dividing the $I_f$ recorded at a given voltage by the cell capacitance. The data were analysed using Clampfit 10.2 (Molecular Devices) and Origin 2024 (Microcal Software Inc.).

**Immunofluorescence of AV node myocytes**

Glass coverslips (13 mm diameter; VWR International GmbH) were coated with poly-L-lysine (PLL) to facilitate cell adhesion. PLL hydrobromide (Sigma-Aldrich, Merck KGaA) was dissolved in deionised H2O to a final concentration of 0.1 mg/ml. Coverslips were sterilised by soaking in 70% ethanol for 1 h, air-dried, then incubated with PLL for 2 h at 37°C, followed by two washes in sterile PBS. AV node myocytes were fixed in 4% paraformaldehyde (PFA) in PBS for 10 min on ice, washed three times for 5 min in PBS–Tween 20 (0.05% v/v), permeabilised in PBS containing Triton X-100 (0.1% v/v) for 10 min and blocked for 1 h in 1% bovine serum albumin (BSA) in PBS. Coverslips were then incubated overnight at 4°C with primary antibodies: rabbit anti-CX3CR1 (1:200, 14-6093-81, Invitrogen; 1:200, SA011F11, Biolegend – qualitatively similar data obtained with both antibodies) and guinea-pig anti-HCN4 (1:250, APC-052-GP, Alomone Labs), diluted in 1% BSA/PBS-T (0.1% Tween 20). After washing, cells were incubated for 3 h with the corresponding secondary antibodies: Alexa Fluor 488 goat anti-rabbit IgG (1:500, A-11008, Thermo Fisher) and Alexa Fluor 647 goat anti-guinea-pig IgG (1:500, A-21450, Thermo Fisher). Coverslips were mounted on glass slides using

ProLong Gold antifade mounting medium with DAPI (Invitrogen). Imaging was performed on a Leica Stellaris 8 STED-FALCON confocal microscope with optimised filters and minimal light exposure. Mouse sinoatrial node and left ventricular myocytes were used as positive and negative controls, respectively, for HCN4 staining. A no-primary-antibody control was included to assess non-specific background staining.

**Mouse tissue preparation and Masson's trichrome staining**

Mouse AV node histology and immunolabelling was performed as described previously.[5] Mouse hearts were excised and immediately flushed with 10 mL of ice-cold Krebs solution to remove residual blood. Dissection was performed to expose the triangle of Koch. Preparations were embedded in optimal cutting temperature (OCT) compound and snap-frozen in −80 °C isopentane. Serial cryosections (10 µm thick) of the AV conduction axis were cut using a Leica CM1860 cryostat equipped with C35 carbon steel blades (Feather) and stored at −80 °C until use. Sections were thawed at room temperature, fixed in 4% PFA for 10 min, and subsequently post-fixed in Bouin's solution overnight. Fixed sections were washed three times in 70% ethanol (10 min each) to remove residual Bouin's solution. For Masson's trichrome staining, sections were stained with Celestine Blue for 5 min, rinsed in three changes of distilled water, and stained with Cole's Alum Haematoxylin for 10 min. After a 15 min wash in tap water (water replaced halfway), slides were stained with Acid Fuchsin for 5 min, washed in distilled water, and treated with phosphomolybdic–phosphotungstic acid for 5 min. Excess reagent was drained before staining with Aniline Blue for 5 min, followed by a brief rinse in distilled water and treatment with 1% acetic acid for 2 min. Dehydration was performed through graded ethanols (70%, 90%, and two changes of 100%; 1–2 min each), followed by clearing in Histo-Clear (National Diagnostics; two changes, 5 min each). Slides were mounted under a fume hood with DPX mounting medium (Sigma-Aldrich), and coverslips were applied carefully to avoid air bubbles.

**Immunofluorescence labelling**

Structures of the AV conduction axis were identified using histology (see above) which was used to guide immunolabelling. Mouse AV conduction axis cryosections were thawed and fixed for 10 minutes in 4% PFA, permeabilised for 10 minutes in PBS containing 0.1% Triton X-100 and blocked for 1 hour in 1% bovine serum albumin (BSA). Washing with PBS-T 0.1% was performed between each step. Primary antibodies were incubated overnight at 4°C, secondary antibodies were incubated for two hours at room temperature. Consistent image acquisition parameters were used for image capture in each experiment. **Single epitope immunofluorescence** Sections were stained with rabbit anti-Cx43 (1:250, C6219, Sigma-Aldrich). Adjacent sections were stained with rabbit anti-HCN4 (1:200, APC-052, Alomone Labs). After washing, Alexa Fluor 488 goat anti-rabbit IgG (1:500, A-11008, Thermo Fisher) was applied. **Double epitope immunofluorescence:** For comparative analysis of macrophage depletion (PLX5622-treated *vs.* control mice), AV conduction axis sections were stained with guinea pig anti-HCN4 (1:200, APC-052-GP, Alomone Labs) and rat anti-CD68 (1:200, ab53444, Abcam), followed by Alexa Fluor 594 goat anti–guinea pig (1:500, A-11076, Thermo Fisher) and Alexa Fluor 488 goat anti-rat (1:500, A21208, Thermo Fisher). **Triple epitope immunofluorescence:** Sections were sequentially stained with guinea pig anti-HCN4 (1:200, APC-052-GP, Alomone Labs) plus Alexa Fluor 647 goat anti–guinea pig (1:500, A-21450, Thermo Fisher); rat anti-CD68 (1:200, ab53444, Abcam) plus Alexa Fluor 555 goat anti-rat (1:500, A-21434, Thermo Fisher); and rabbit anti-Cx43 (1:200, C6219, Sigma-Aldrich) plus Alexa Fluor 488 goat anti-rabbit (1:500, A-11008, Thermo Fisher). **Imaging:** After immunolabelling, all slides were mounted with ProLong Gold antifade mountant containing DAPI (Invitrogen). Single- and double-label images and endogenous GFP in *Cx$_3$cr1$^{GFP/+}$* mice were captured on an Axioimager fluorescence microscope (Zeiss); triple-label images were acquired on an LSM-780 inverted confocal laser-scanning microscope (Zeiss).

The compact node was defined by its position adjacent to the AV septal musculature and consisted of a knot of small, densely packed HCN4-positive myocytes within the central fibrous body. The inferior nodal extension was identified histologically as an extension of HCN4-positive cells projecting from the base of the crista terminalis region to the compact node. More distal sections revealed the penetrating bundle, a direct continuation of the compact node, histologically recognisable as a well-organised bundle of small, interlacing HCN4-positive myocytes encased in the central fibrous body at the junction of the interatrial and interventricular septa. This structure continued distally as the proximal His bundle, which traversed the insulating connective tissue before bifurcating into the left and right bundle branches and eventually terminating in Purkinje fibres embedded within the subendocardial connective tissue of the ventricular septum.

**Co-localisation quantification**
Mouse AV node sections were imaged using a 20× dry objective, and human AVN sections were imaged using a 10× dry objective. For each sample, images were collected as Z-stacks (5 optical sections) with a Z-step size of 3–5 µm, and maximum intensity projections were generated for analysis. Co-localization analysis was performed in Fiji/ImageJ (version 1.54p) using the JAcoP plugin (*Just Another Co-localization Plugin*). Fluorescence thresholds were standardized by calculating the mean optimal threshold across all images within each experiment. Co-localization was quantified using Manders' overlap coefficients (M1 and M2), which measure the fraction of one fluorophore's signal overlapping with the other. Analyses were performed for HCN4 with Cx43 and Cx43 with CD68 within HCN4-positive regions of the mouse AV node, and for Cx43 with CD68 in the human AV node. Data are presented as mean ± standard deviation (n = 3–4 images per slide from 15 slides per sample from 3 human hearts per group).

**Human tissue preparation and Masson's Trichrome staining**
Human atrioventricular (AV) nodes were dissected and embedded in optimal cutting temperature compound (OCT) and stored at −80 °C. Serial cryosections (10 µm thick) were obtained using a Leica CM1860 cryostat equipped with C35 carbon steel blades (Feather) and stored at −80 °C until use. For histological assessment, sections were thawed for 10 min at room temperature and fixed overnight in Bouin's solution (Sigma-Aldrich) to enhance connective tissue staining contrast. Excess Bouin's solution was removed by washing three times in 70% ethanol. Sections were then incubated sequentially in Weigert's iron haematoxylin (10 min), Biebrich scarlet–acid fuchsin (5 min), phosphomolybdic–phosphotungstic acid (5 min), and aniline blue (5 min), with brief washes in distilled water between steps. Sections were rinsed briefly in 1% acetic acid, dehydrated through graded ethanol (70%, 90%, and two changes of 100%), cleared twice in Histo-Clear (National Diagnostics; 5 min each), and mounted in DPX mounting medium (Sigma-Aldrich).

**Human tissue immunofluorescence labelling**
Frozen human AV node sections were fixed for 10 min in 4% paraformaldehyde (PFA) (Thermo Fisher Scientific) at room temperature. Sections were permeabilized for 10 min in PBS containing 0.1% Triton X-100, then blocked for 1 h in buffer containing 2.5% bovine serum albumin (BSA; Sigma-Aldrich), 5% goat serum (Thermo Fisher Scientific), and 0.1% Triton X-100 in PBS-T (PBS + 0.1% Tween-20).
To reduce autofluorescence, sections were treated with 3% hydrogen peroxide (Thermo Fisher Scientific) for 10 min, followed by PBS-T washes. Primary antibody incubation was performed overnight at 4 °C using:

- Rabbit anti-Connexin 43 (Cx43), 1:500 (C6219, Sigma-Aldrich)
- Mouse anti-CD68, 1:100 (14-0688-82, Thermo Fisher Scientific)

After washing, sections were incubated for 1 h at room temperature with:

- Alexa Fluor 488 goat anti-rabbit IgG, 1:500 (A11008, Thermo Fisher Scientific)
- Alexa Fluor 594 goat anti-mouse IgG, 1:500 (A11032, Thermo Fisher Scientific)

Slides were washed again in PBS-T, mounted with Vectashield Antifade Mounting Medium containing DAPI (Vector Laboratories), and imaged using a Leica STELLARIS 8 confocal microscope equipped with 405, 488, and 594 nm laser lines. Identical acquisition parameters were used for all samples.

**Macrophage depletion and ex vivo electrophysiological study**
Macrophage depletion was achieved using the colony-stimulating factor 1 receptor (CSF1R) inhibitor, PLX5622. Mice were fed a PLX5622-supplemented diet (1200 ppm) formulated into pellets (Research Diets Inc.) for 21 days, while controls received a standard rodent diet. Anaesthetised electrocardiograms (ECGs) were recorded before and after treatment as described previously.[5,6] At the end of the treatment period, hearts were excised and mounted on a Langendorff perfusion apparatus for ex vivo electrophysiological study, perfused with Krebs buffer bubbled with carbogen.

Contact electrodes were positioned at the left atrial appendage and apex, with a pacing electrode placed at the right atrial appendage (Mapping Lab system). After a 10-minute stabilisation period, a pseudo-ECG was recorded. Hearts were paced at multiple cycle lengths to determine PR interval. The Wenckebach cycle length was assessed by atrial pacing with decremental pulse trains, reducing the cycle length by 1 ms per step until 1:1 atrioventricular conduction was lost. The AV nodal effective refractory period was determined using an S1S2 protocol consisting of eight S1 stimuli at 120 ms, followed by an S2 stimulus decrementing by 1 ms until AV conduction failed (i.e., a dropped QRS complex).[5,7,8]

**Macrophage quantification**
Following electrophysiological study, the triangle of Koch was micro-dissected, weighed and placed in 4°C Krebs solution. Tissue was digested for one hour in; collagenase I (450u/ml, Worthington Biochemical), collagenase XI (125u/ml, Sigma Aldrich), DNAse from bovine pancreas (60u/ml, Merck), Hepes buffer (1 molar 20 µl/ml, Sigma Aldrich) and hyaluronidase from bovine testes (60u/ml, Sigma Aldrich). During this time the samples were kept at 37.5°C on a heat block in 2 ml Eppendorff tubes which were inverted, and briefly vortexed at low speed, every 3 minutes. Following digestion, the suspension was filtered through a 40µm cell strainer. This protocol was adapted from Ruibing *et al.*[9] Cells were stained with Brilliant Violet 605™ anti-mouse Ly-6C Antibody [Clone: HK1.4] PE anti-mouse F4/80 Antibody [Clone: BM8], APC anti-mouse Ly-6G Antibody [Clone: 1A8], Brilliant Violet 421™ anti-mouse/human CD11b Antibody [Clone: M1/70], FITC anti-mouse CD45 Antibody (All Biolegend, 1:100 and Zombie NIR™ Fixable Viability Kit, 1:2500). Cells were fixed in 2% paraformaldehyde for 10 minutes then washed and analysed on a BD FACS Canto II flow cytometer. Data was analysed using FlowJo™ v11 Software.

**Clearing and staining of mouse AV node**
AV node tissue was permeabilised in phosphate buffered saline containing triton-X 100 (PBST (0.3% triton x-100)) at room temperature overnight to permeabilize the tissue. Next the tissue was incubated in blocking buffer PBS +1% BSA, overnight on a shaker at 4°C. Primary antibody was prepared in blocking buffer and the tissue was incubated in the primary antibody cocktail (containing guineapig anti-mouse HCN4 (APC-052-GP Alomone Labs); rat anti-mouse CD68 1:200 (ab53444, Abcam); rabbit anti-mouse Cx43 (C6219, Sigma-Aldrich). – all 1:200) for 48h with gentle agitation at 4°C. The primary antibody was washed in 3 x PBST (PBST (0.01% triton x-100)) and incubated for 24h in secondary antibody (containing goat anti-guinea pig 647 A-21450, ThermoFisher; goat anti-rat 555, A-21434, ThermoFisher, goat anti-rabbit Alexa488, A-11008, ThermoFisher – all 1:500, and DAPI) at 4°C for 24 hours. The secondary antibody was washed off in 3 x PBST (PBST (0.01% triton x-100)) and then 2 x PBS. Next the stained tissue was embedded in 2% low melting point agarose (diluted in MilliQ water). Tissue block was allowed to set in the fridge overnight and then dehydrated in series of ethanol – from 25%, 50%, 70%, 90% to 100%. 1.5 hours for each step from 25-90% and 2 hours for 100% ethanol. The dehydrated agarose-tissue block was incubated in BABB (Benzyl alcohol, benzyl benzoate; 1:2) at room temperature overnight to dissolve tissue lipids and the cleared tissue was stored and imaged in ethyl cinnamate and imaged on an SP5 confocal microscope using a µ-Plate 24 Well IBIDI (Thistle Scientific).

**Imaging**
Cleared AV node tissue was imaged using a Leica SP5 MP/FLIM inverted microscope (Leica, Wetzlar, Germany). DAPI was excited with a 405nm diode laser, Cx43 was excited using a 488nm argon laser, CD68 were excited using a 543 nm HeNe laser, while HCN4 was excited using a 633nm HeNe laser. A tile scan was performed with images acquired using 1024 x 1024 pixel format with a z-stack step size of 0.5 micron. Images were acquired with a 16bit imaging depth. Maximum projection images were generated, and 3D visualisation and export were performed on Imaris v9.6.0 (Oxford Instruments).

**Statistics**
Statistical analysis was performed on GraphPad Prism software (10.2.3). Normality was assessed using Shapiro-Wilk's test. For normally distributed data, Welch's t test was used throughout. If

normality was not confirmed, a Mann-Whitney test was used. N numbers for all experiments are stated in the figure legends. All bar graphs denote mean±standard deviation.

### SUPPLEMENTAL FIGURE LEGENDS

**Figure S1.** AV node myocyte membrane potential during pacing at 550 beats/min when coupled to Type 0 (A), Type 1 (C), Type 2 (E) or Type 1+2 (G) macrophage models at Ggap = 0, 3 or 7 nS. For each condition, the corresponding macrophage membrane potential is also shown. (B,D,F,H) Myocyte ionic currents for the same macrophage phenotypes at Ggap = 3 nS, including background current current ($I_b$), inward rectifier $K^+$ current $I_{K,1}$ ($I_{Kir2.1}$), ultrarapid delayed rectifier $K^+$ current $I_{K,ur}$ ($I_{Kv1.5}$), gap junction current from the perspective of the macrophage ($I_{GapM\phi}$) and the same gap junction current but from the perspective of the myocyte ($I_{GapMyo}$). Positive IGap_Myo denotes current leaving the myocyte (reverse sign for IGap_MΦ). The Y axes are interrupted for clarity.

**Figure S2. Coupling AV node myocytes to macrophages depolarises the myocyte resting potential, prolongs action potential duration and impairs conduction.** (A,B) Relationship between the myocyte resting potential (A) and $APD_{90}$ (B) and the coupling conductance between the myocyte and four macrophage subtypes (stimulation rate, 550 beats/min). (C, D) Relationship between the maximum diastolic potential (MDP; C) and rate of spontaneous activity (D) of the myocyte and the coupling conductance between the myocyte and the four macrophage subtypes. (E) 30 s simulation of membrane potential of myocyte 20 (100 myocyte strand; Type 0 macrophage; 350 beats/min) in the absence (left; coupling conductance 0 nS) and presence (right; coupling conductance 3 nS) of macrophage coupling, illustrating the persistence of 2:1 block. (F) Relationship between conduction velocity and coupling conductance for coupling between AV node myocytes and the four types of macrophage (100 myocyte strand; 350 beats/min).

**Figure S3. CX3CR1 is expressed by nodal myocytes.** (A) AV node myocytes isolated from the Triangle of Koch of a wild-type mouse (region of interest shown in Fig. 2A) double immunolabelled for HCN4 and CD68 (top) or HCN4 and CX3CR1 (bottom) and stained with DAPI to highlight nuclei. Similar data were obtained from 24 myocytes isolated from n=3 mice. (B) Left, the penetrating bundle of a $Cx_3cr1^{GFP/+}$ mouse identified by histology. The section was immunolabelled for HCN4 and stained with DAPI. Right, high magnification images (from the boxed region on the left) of the DAPI, HCN4 and GFP signals as well as a composite image of all three signals. White arrows on the composite image, areas of signal co-localisation of GFP (*Cx₃cr1* marker) and HCN4. (C) Three representative AV node myocytes isolated from the Triangle of Koch of the $Cx_3cr1^{GFP/+}$ mouse immunolabelled for HCN4 and stained for DAPI. For each myocyte, the images show the GFP signal ($Cx_3cr1$ marker), the HCN4 signal, and the composite image together with the DAPI signal. (D) Mean expression levels of $Cx_3cr1$ in component cells of the mouse sinoatrial node determined using single nucleus RNA sequencing (GEO Series accession number GSE130710).

**Figure S4. Expression of Cx43 and HCN4 throughout the AV conduction axis of the mouse.** Numbered long axis serial cryosections through the AV conduction axis of a mouse were stained with Masson's trichrome and immunolabelled for Cx43 and HCN4. For each section, a low magnification image of Masson's trichrome staining is shown at the top, a high magnification image of Cx43 immunolabelling is shown in the middle, and a high magnification image of HCN4 immunolabelling is the shown at the bottom. The dotted lines identify structures of the AV conduction axis: the Bundle of His (sections 1-4), the penetrating bundle (sections 5-10), the compact AV node (sections 11-13) and the inferior nodal extension (sections 14 and 15). Scale bar = 200 µm.

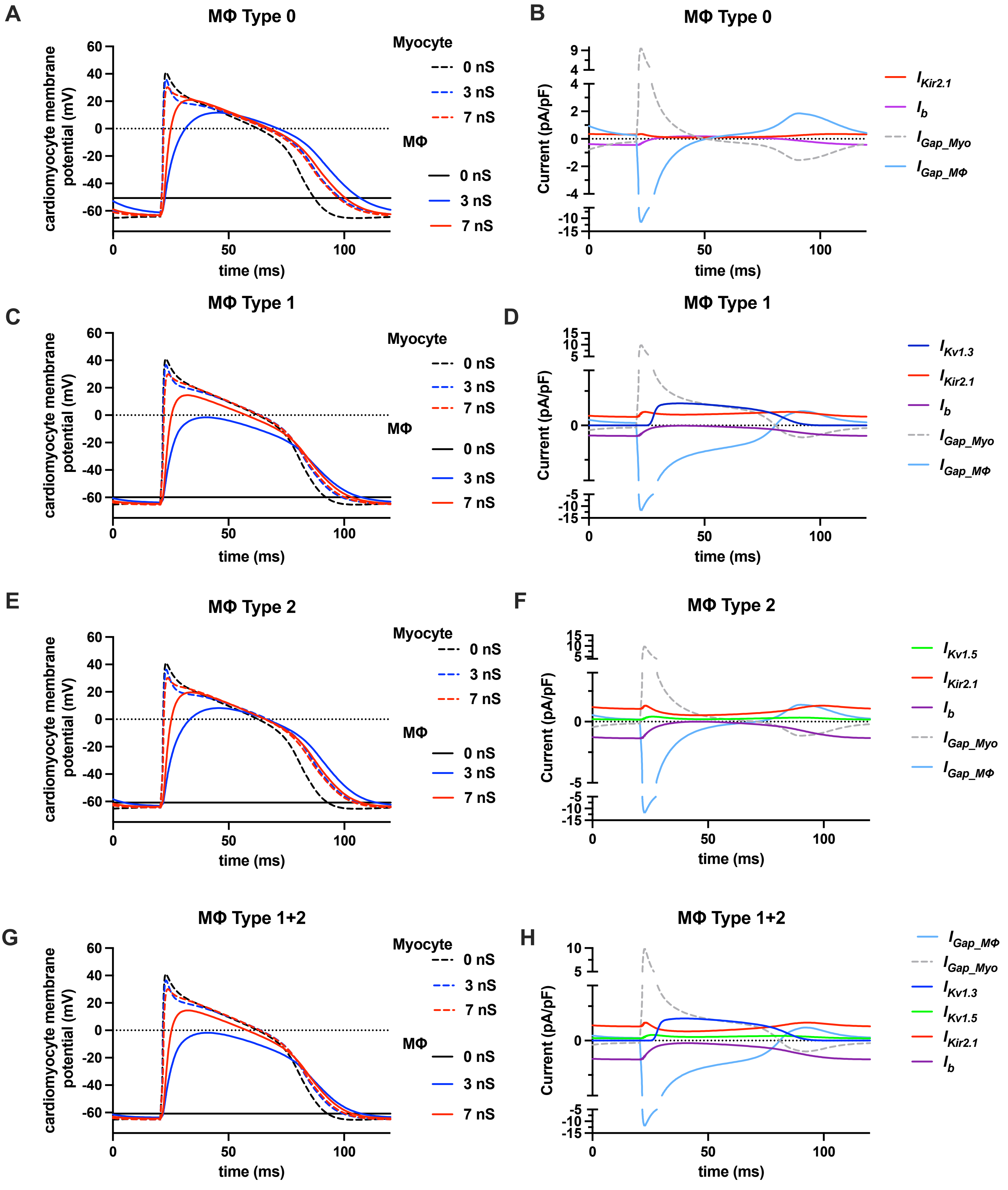
A
MΦ Type 0
cardiomyocyte membrane potential (mV)
time (ms)
Myocyte
0 nS
3 nS
7 nS
MΦ
0 nS
3 nS
7 nS
B
MΦ Type 0
Current (pA/pF)
time (ms)
$I_{Kir2.1}$
$I_b$
$I_{Gap\_Myo}$
$I_{Gap\_MΦ}$
C
MΦ Type 1
cardiomyocyte membrane potential (mV)
time (ms)
Myocyte
0 nS
3 nS
7 nS
MΦ
0 nS
3 nS
7 nS
D
MΦ Type 1
Current (pA/pF)
time (ms)
$I_{Kv1.3}$
$I_{Kir2.1}$
$I_b$
$I_{Gap\_Myo}$
$I_{Gap\_MΦ}$
E
MΦ Type 2
cardiomyocyte membrane potential (mV)
time (ms)
Myocyte
0 nS
3 nS
7 nS
MΦ
0 nS
3 nS
7 nS
F
MΦ Type 2
Current (pA/pF)
time (ms)
$I_{Kv1.5}$
$I_{Kir2.1}$
$I_b$
$I_{Gap\_Myo}$
$I_{Gap\_MΦ}$
G
MΦ Type 1+2
cardiomyocyte membrane potential (mV)
time (ms)
Myocyte
0 nS
3 nS
7 nS
MΦ
0 nS
3 nS
7 nS
H
MΦ Type 1+2
Current (pA/pF)
time (ms)
$I_{Gap\_MΦ}$
$I_{Gap\_Myo}$
$I_{Kv1.3}$
$I_{Kv1.5}$
$I_{Kir2.1}$
$I_b$

Fig. S2

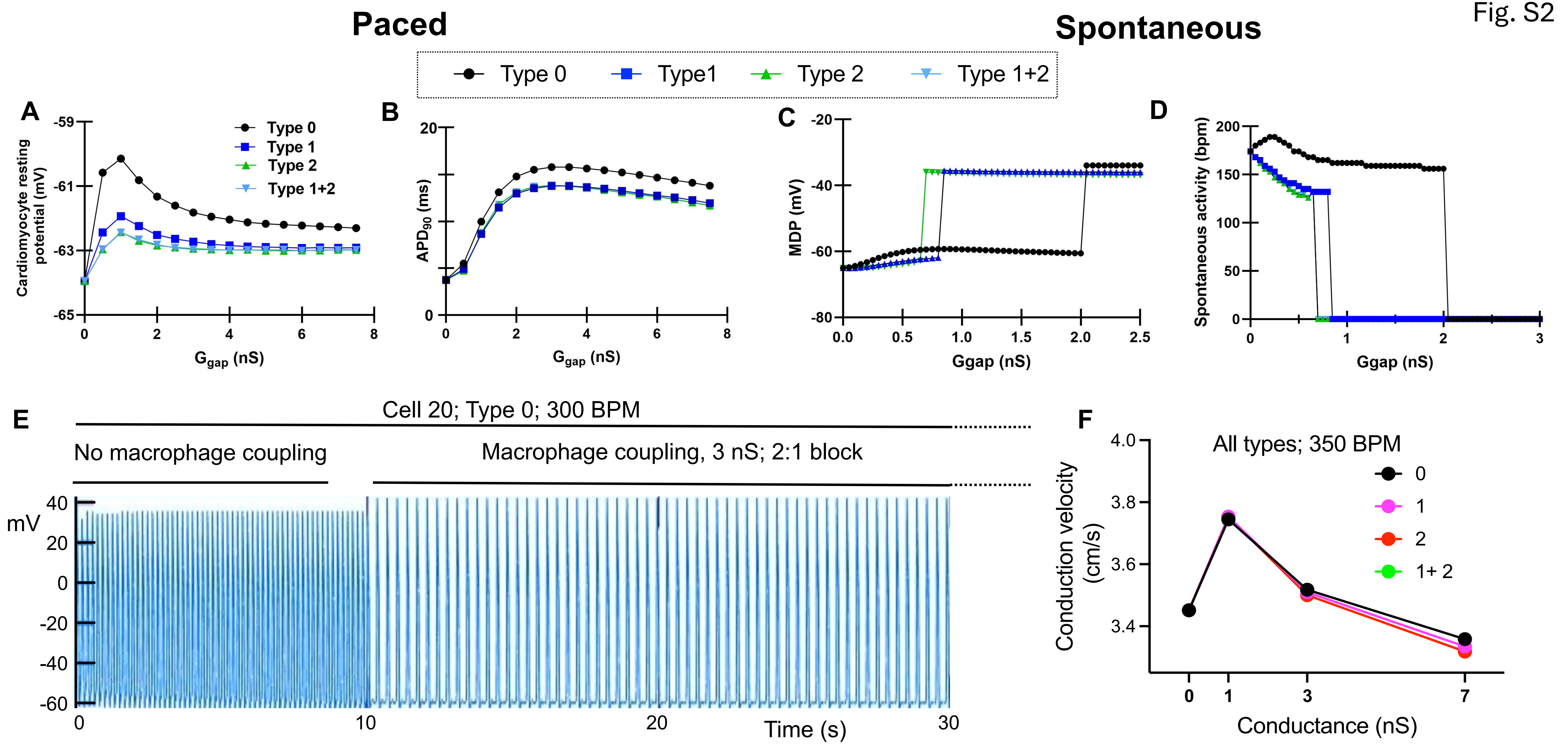

Paced
Spontaneous
Type 0
Type1
Type 2
Type 1+2
A
Cardiomyocyte resting potential (mV)
$G_{gap}$ (nS)
Type 0
Type 1
Type 2
Type 1+2
B
$APD_{90}$ (ms)
$G_{gap}$ (nS)
C
MDP (mV)
Ggap (nS)
D
Spontaneous activity (bpm)
Ggap (nS)
E
Cell 20; Type 0; 300 BPM
No macrophage coupling
Macrophage coupling, 3 nS; 2:1 block
mV
Time (s)
F
All types; 350 BPM
Conduction velocity (cm/s)
Conductance (nS)
0
1
2
1+ 2

Fig. S3

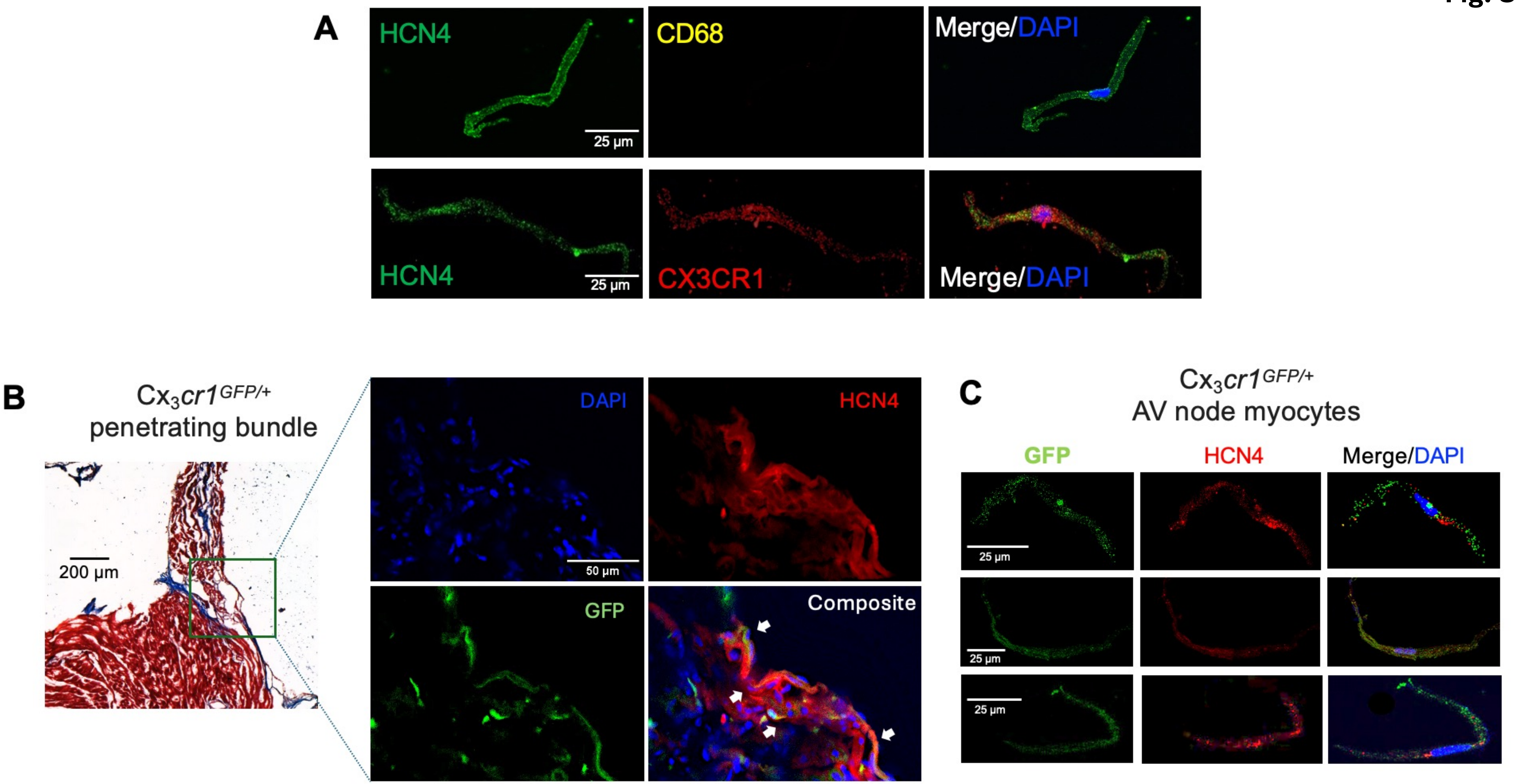
A
HCN4
25 µm
CD68
Merge/DAPI
HCN4
25 µm
CX3CR1
Merge/DAPI
B
$Cx_3cr1^{GFP/+}$
penetrating bundle
200 µm
DAPI
50 µm
HCN4
GFP
Composite
C
$Cx_3cr1^{GFP/+}$
AV node myocytes
GFP
HCN4
Merge/DAPI
25 µm
25 µm
25 µm

Fig. S4

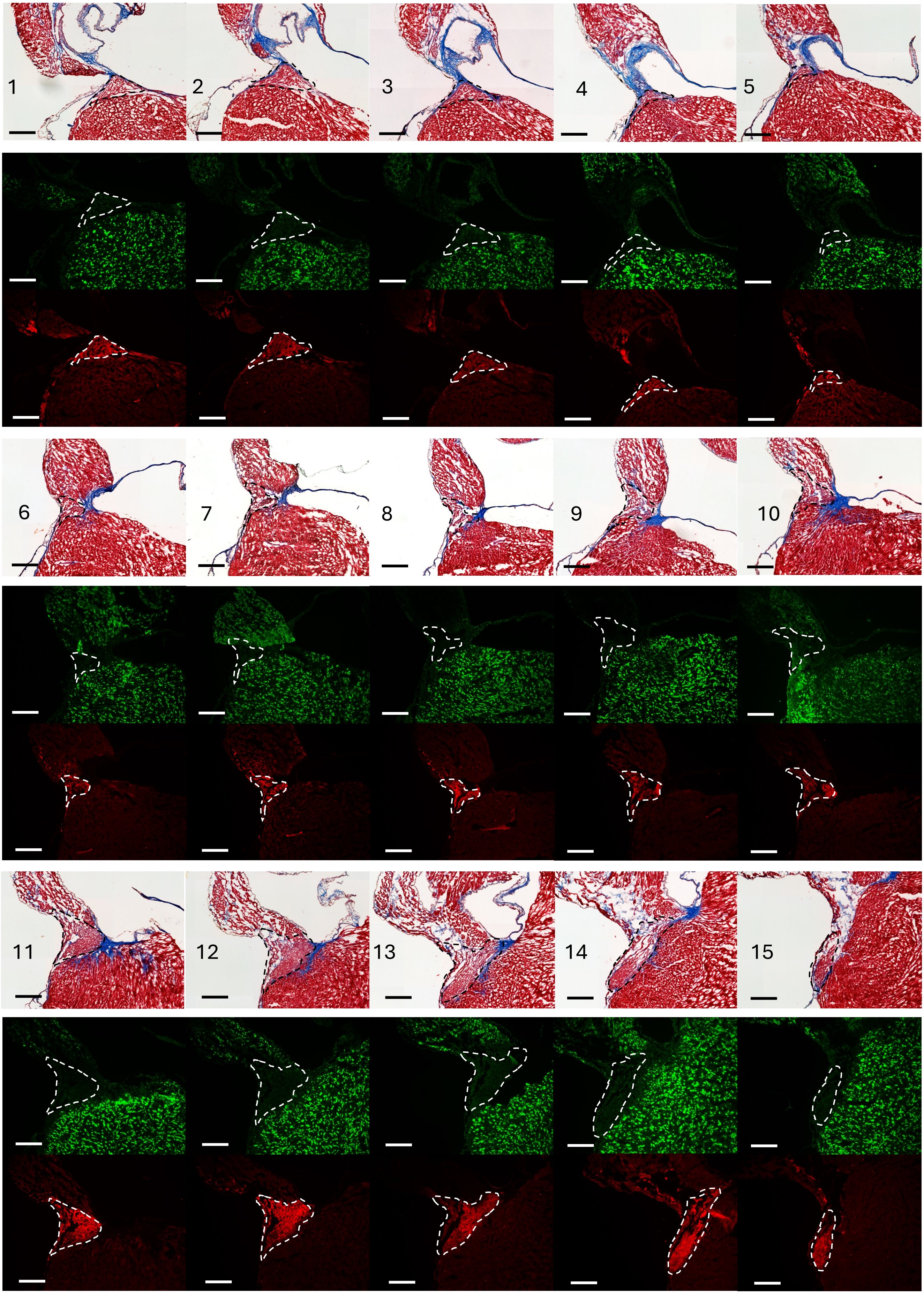
